# ACSF2: A MEDIUM-CHAIN ACYL-CoA SYNTHETASE WITH A POTENTIAL ROLE IN NEURONAL DIFFERENTIATION

**DOI:** 10.1101/2022.03.28.486105

**Authors:** Dony Maiguel, Masashi Morita, Zhengtong Pei, Zhenzhen Jia, Paul A. Watkins

## Abstract

By activating fatty acids to their CoA derivatives, acyl-CoA synthetases (ACS) play an essential role in fatty acid metabolism. We previously identified ACSF2 as an ACS that was phylogenetically distinct from known families of short-chain, medium-chain, long-chain, very long-chain, and bubblegum ACSs. Functionally, ACSF2 preferentially activated medium-chain fatty acids. In this work, we provide further characterization of this unique ACS. ACSF2 mRNA expression was found in most tissues, although immunohistochemical analysis revealed differences in protein expression between various cell types in each tissue. Endogenous ACSF2 was found in the Golgi region in Neuro2a and P19 cells, and disruption of the Golgi in Neuro2a cells with nocodazole disrupted ACSF2 localization. In contrast, MA-10, HepG2, and skin fibroblasts had a mitochondrial ACSF2 localization. ACSF2 activated saturated fatty acids containing 6 to 10 carbons when overexpressed in COS-1 cells. The Km_app_ for C10:0 was 24.4 μM and Vmax_app_ was 385 nmol/20min/mg COS cell protein. Knockdown by RNA interference revealed that ACSF2 was responsible for most of the medium-chain ACS activity in Neuro2a cells. A lysine residue critical for activity in bacterial short-chain ACSs was found in ACSF2, and mutation of this residue to alanine abolished enzyme activity. Neurite outgrowth results when Neuro2a cells are induced to differentiate with retinoic acid, and ACSF2 migrated to nodes and points of neurite-neurite contact along with the presynaptic marker, synaptophysin. ACSF2-deficient Neuro2a cells showed significantly blunted neurite outgrowth in response to retinoic acid. These results suggest that this medium-chain ACS may play an important role in neuronal development.

## INTRODUCTION

Utilization of fatty acids (FA)^1^ is essential for normal cellular function. FAs are required for a variety of processes including formation of membrane phospholipids and sphingolipids, energy storage and metabolism, neurometabolism, and protein covalent modification. FA uptake by cells is followed by activation of the FA by thioesterification to coenzyme A (CoA), a reaction catalyzed by acyl-CoA synthetases (ACSs) [1]. Many early studies revealed that tissues contained distinct ACS for activating short-, medium-, long-, and very long-chain FAs, and it now well-established that for each FA chain length category, multiple ACSs exist [1,2].

Very long-chain fatty acids (VLCFA, containing 22 or more carbons) are particularly abundant in brain [3]. Disturbed VLCFA metabolism and increased tissue levels of VLCFA are seen in several human peroxisomal diseases. Neurodegeneration is a common feature of these disorders, which include X-linked adrenoleukodystrophy, deficiency of either peroxisomal acyl-CoA oxidase or D-bifunctional protein, and the peroxisome biogenesis disorders [4-6]. In order to understand the mechanism of VLCFA accumulation in these diseases, we sought to identify and characterize ACSs capable of activating VLCFA. Using bioinformatic approaches, we concluded that there are 26 ACSs in the human, rat and mouse genomes [2]. While most candidate ACSs segregated into phylogenetically distinct families of short-chain (ACSS), medium-chain (ACSM), long-chain (ACSL), very long-chain (SLC27A), and “bubblegum” (ACSBG) ACSs, four proteins did not. These were given the designation ACSF (ACS Family) members 1-4 [7]. Because their structures do not provide clues regarding chain-length preference, they must be characterized independently.

We cloned ACSF2 and ACSF3 cDNA, overexpressed the proteins in COS-1 cells, and demonstrated that both had ACS enzyme activity [2]. While ACSF3 showed a weak ability to activate a 24-carbon very long-chain FA, it was subsequently characterized as malonyl-CoA and methylmalonyl-CoA synthetase [8,9]. Mutations in ACSF3 are causative of combined malonic and methylmalonic aciduria [9]. In contrast, our preliminary analysis of ACSF2 suggested it was a medium-chain ACS. Here, we verify that ACSF2 is indeed a medium-chain ACS with a broad tissue distribution. We demonstrate that an evolutionarily conserved lysine residue near the C-terminus is critical for the catalytic activity of ACSF2. Furthermore, we show that the subcellular location of ACSF2 differs in different cell types. Using a mouse neuroblastoma cell model, we demonstrate that this protein is required for neurite outgrowth and, perhaps, synapse formation.

## EXPERIMENTAL PROCEDURES

### Materials and general methods

[1-^14^C]lauric acid (C12:0), [1-^14^C]palmitic acid (C16:0), [1-^14^C]arachidonic acid (C20:4), [1-^14^C]docosahexaenoic acid (C22:6), and [1-^14^C]lignoceric acid (C24:0) were obtained from Moravek, Inc. (Brea, CA). [1-^14^C]octanoic acid (C8:0), [1-^14^C]decanoic acid (C10:0), [1-^14^C]myristic acid (C14:0), [1-^14^C]oleic acid (C18:1) and [1-^14^C]linoleic acid (C18:2) were obtained from American Radiolabeled Chemicals (St. Louis, MO). [1-^14^C]stearic acid was obtained from Amersham Biosciences. [1-^14^C]hexanoic acid (C6:0) was from ViTrax (Placentia, CA). Unlabeled fatty acids were from either Sigma Chemical or Cayman Chemical. Protein was measured by the method of Lowry et al. [10]. General conditions for PCR were as previously reported [11]. DNA sequencing was performed at the Johns Hopkins University Dept. of Biological Chemistry Biosynthesis and Sequencing Facility using the fluorescent di-deoxy terminator method of cycle sequencing on an Applied Biosystems Inc. 377 automated DNA sequencer, following ABI protocols. COS-1 cells were transfected by electroporation as described previously [11]. Sources of commercial antibodies were: synaptophysin (mouse monoclonal; Sigma), Golgin 97 (mouse monoclonal; Molecular Probes), GM130 (mouse monoclonal; BD Bioscience), ATP synthase (mouse monoclonal; Chemicon), manganese superoxide dismutase (rabbit polyclonal; Stressgen), donkey anti-rabbit IgG FITC conjugate and donkey anti-mouse IgG rhodamine conjugate (Jackson ImmunoResearch), goat anti-rabbit IgG horseradish peroxidase conjugate (Santa Cruz), goat anti-mouse IgG Alexa fluor 680 conjugate (Molecular Probes), and goat anti-rabbit IgG IR dye 800 conjugate (Rockland). Kinetic parameters were obtained using GraphPad Prism 3.03. Statistical significance was calculated using Student’s t-test.

### Animals and care

Wild type 129SvEv mice were obtained from Taconic, Inc. (Germantown, NY) and were approximately three months of age. Mice were housed in facilities of the Johns Hopkins University School of Medicine under controlled conditions, between 22°C and 27°C, on a 12-h light/dark cycle, with food and water *ad libitum*. Procedures involving animals and their care were conducted in conformity with institutional guidelines that are in compliance with national and international laws and policies (EEC Council Directive 86/609,OJ L 358, 1 DEC.12,1987; NIH Guide for the Care and Use of Laboratory Animals, U.S. National Research Council, 1996).

### Cloning of full-length mouse ACSF2 cDNA and site-directed mutagenesis

Full-length cDNA encoding mouse brain ACSF2 was cloned as previously described [2]. The overlap extension method of Ho et al. [12] was used to mutate lysine 599 of mACSF2 to alanine, as described previously [13]. Briefly, full-length mACSF2 cDNA was used as template to amplify by PCR two overlapping fragments so that each incorporates the desired mutation. Forward primer 5’-AAATTTGGATCCAGAGCCATGGCTGTCTATCAC-3’ (mACSF2-f) and reverse primer 5’-GAATTTCTGGAT<u>GGC</u>TCCTGAGATGGT-3’ were used to amplify a fragment encoding amino acids 1-599. Forward primer 5’-ACCATCTCAGGA<u>GCC</u>ATCCAGAAATTC-3 and reverse primer 5’-TTTAAACTCGAGACCCTCCTTTGCTTCACAGT-3’ (mACSF2-r)were used to amplify the fragment encoding amino acids 599-615. The underlined GCC (A) codon replaced the wild-type AAG (K) codon. The PCR products were purified, combined, and used as template in a second round of PCR with primers mACSF2-f and mACSF2-r. The final PCR product was cloned into the BamH1 and Xho1 sites of pcDNA3, and re-sequenced. Except for the desired mutation, the sequence was identical to that of the full-length wild-type cDNA.

### Cell Culture conditions

Mouse neuroblastoma Neuro2a cells, monkey kidney COS-1 cells, and human hepatoma HepG2 cells were from American Type Culture Collection (Manassas, VA) and grown in DMEM media (Invitrogen, CA) containing 10% fetal bovine serum (Gemini Bioproducts, CA). Fibroblasts were generated from newborn mouse skin and cultured in the same medium. MA-10 cells [14] were a gift from Dr. Mario Ascoli and were cultured as described previously [15]. P19 mouse teratocarcinoma cells were from American Type Culture Collection and were grown in α-MEM supplemented with 7.5% newborn bovine serum + 2.5% fetal bovine serum. All cells were grown in an incubator at 37°C in a 5% CO_2_ atmosphere.

### RNA interference studies

For transient knockdown of ACSF2, a double-stranded small inhibitory RNA (siRNA) construct was prepared as described previously [16] using the following oligonucleotides: 5’-AAGCGAGCCATGGCTGTCTATCCTGTCTC-3’ and 5’-AAATAGACAGCCATGGCTCGCCCTGTCTC-3’. This constructs targets bp 8-27 of the mouse ACSF2 coding sequence. Transfection of siRNA was done using Ambion siPORT Amine (Austin, TX) according to manufacturer’s instructions with the exception that the cells were incubated with siRNA in serum free media for ten hours before the addition of fresh media containing fetal bovine serum. To produce a stable mACSF2 knockdown cell line, an shRNA-generating sequence equivalent to the above siRNA sequence was inserted into the pSilencer 4.1 CMV-Hygro vector (Ambion, Austin TX). The plasmid was transfected into Neuro2a cells and clonal lines selected by incubation with Hygromycin (300 mg/ml, Roche (Nutley, NJ)). After two weeks, antibiotic-resistant colonies were selected, expanded, and checked for mACSF2 knockdown by indirect immunofluorescence and Western blot. A control cell line was similarly generated using a random shRNA sequence inserted into the pSilencer 4.1 CMV-Hygro vector.

### Nocodazole treatment

Neuro2a cells were grown on glass coverslips and incubated with 20 μM nocodazole (Calbiochem, San Diego CA). Coverslips were removed at zero time and 2 hours and were fixed for 10 min using 3% formaldehyde in PBS prior to indirect immunofluorescence analysis as described below. Hoechst staining was used to visualize nuclei.

### Retinoic acid treatment

Stable mACSF2 knockdown cells and control cells were grown on glass cover slips and treated with 40 μM all-*trans*-retinoic acid (Calbiochem, San Diego CA). Forty eight hours post transfection cells were fixed for 20 min with a solution of 3% formaldehyde in PBS. Images were obtained from Nikon T1000 eclipse microscope using 200x magnification. Using Adobe Photoshop, three concentric circles were superimposed on the images and the number of cells whose neurites crossed one, two or three circles was counted. As criteria, only cells that formed clusters of three cells or less were counted, and only the number of circles the longest neurite crossed was counted for each cell. The diameters of the three circles were calculated to be 22, 48, and 73 μm. Data from three independent treatments of knockdown and control cell lines were analyzed. For each treatment, twenty images were obtained. A total of approximately 14,000 cells were examined.

### Northern Blot Analysis

Total RNA was prepared from freshly harvested mouse tissues using the Trizol reagent (Invitrogen). Prenatal and newborn mouse brain RNA was kindly provided by Dr. Phyllis Faust (Columbia Univ.). A 956 bp cDNA probe was prepared by PCR amplification using forward primer 5’-TCTTCTGAGCATTGGCCTCC-3’ and reverse primer 5’-CCACACTACCAGCCTTCTGC-3’, and full-length *ACSF2* cDNA as template. Northern blot analysis was done essentially as described previously [15]. For control, a 528 bp GAPDH probe was used.

### ACS activity assay

Activation of [1-^14^C]fatty acids to their CoA derivatives was assayed as previously described [15]. In standard assays, the reaction mix contained 40 mM Tris (pH 7.5), 10 mM ATP, 10 mM MgCl_2_, 0.2 mM CoA, 0.2 mM DTT and 20 μM labeled fatty acid. The fatty acyl-CoA product was separated from the substrate either by the method of Dole [17] or the method described by Fujino et al. [18]. For assessment of pH optimum, potassium phosphate buffer was used instead of Tris. In some experiments, the fatty acid concentration was 400 μM as indicated in the figure legends.

### FA β-oxidation and incorporation into complex lipids

Degradation of [1-14C]C10:0 to water-soluble products by β-oxidation was measured as previously described [19], except that the final FA concentration in the assay was 0.4 mM. To investigate the incorporation of [1-14C]C10:0 into complex lipids, cells were harvested by gentle trypsinization, washed with PBS, and resuspended in 0.25 M sucrose containing 10 mM Tris, pH 8.0, and 1 mM EDTA as for β-oxidation studies. Labeled FA (0.4 mM final concentration) was solubilized with α-cyclodextrin (10 mg/ml in 10 mM Tris, pH 8.0), also as for β-oxidation studies. Reaction mixes contained (final concentrations): Tris, pH 7.5, (40 mM), ATP (10 mM), MgCl_2_ (10 mM), CoA (0.2 mM), and α-glycerophosphate (1 mM) in a total volume of 250 μl, for 2 h at 37°C. Reactions were terminated by the addition of chloroform/methanol (2:1) containing 5 mM HCl, and extraction was carried out according to the method of Folch et al. [20]. The organic phase was dried under a stream of nitrogen and solubilized with 50 μl chloroform. Twenty μl aliquots were applied to Whatman Lk6D silica gel high performance thin layer plates. Plates were pre-washed with chloroform/methanol (1:1), and for phospholipid analysis only, plates were subsequently wetted with 2.3% boric acid in ethanol and drained 5 min. All plates were dried for 15 min at 100°C. Solvent systems used were hexane/diethyl ether/acetic acid (80:20:1) for neutral lipids and chloroform/ethanol/water/triethylamine (30:35:7:35) for phospholipids. Labeled lipids were detected by phosphorimager analysis (Fuji-BAS 2500) and identified by comparison to authentic standards. Neutral lipid standards were detected by exposure to iodine vapor, and phospholipid standards were detected under ultraviolet light after spraying the plates with primuline (0.005% (w/v) in acetone/water (4:1)).

### ACSF2 antibody production and purification

Antibody to ACSF2 was raised against a fusion protein containing maltose-binding protein and the C-terminus of ACSF2. Full-length ACSF2 in pcDNA3 was used as template to amplify by PCR an ∼0.5 kb fragment encoding the C-terminal 165 amino acids with forward oligonucleotide primer 5’-AAATTT<u>GAATTC</u>GAGCTGACCAACCTGAACGTG-3’ (which incorporates an EcoRI site, underlined) and reverse primer 5’-AGATGGCTGGCAACTAGAAGG-3’ (which hybridizes beyond the XbaI site of pcDNA3). The PCR product was ligated into the EcoRI and XbaI sites of the pMAL-c2 vector (New England Biolabs, Ipswich, MA). The resulting plasmid was transduced into *E. coli* strain DH10b. Protein expression was induced for 3 h at 30°C with 0.4 mM isopropyl-β-D-thiogalactoside. Washed cells were disrupted by sonication in 20 mM Tris, pH 7.5, 0.2 M NaCl, 1 mM EDTA, 1mM DTT, and 0.5 mM PMSF and centrifuged at 20,000 x g. The fusion protein was purified from the supernatant using amylose resin (New England Biolabs, Ipswich, MA), following the manufacturer’s protocol. Polyclonal antiserum to the fusion protein was raised in a rabbit by Cocalico Biologicals (Reamstown, PA). For affinity purification, 0.5 mg of fusion protein was subjected to preparative SDS-PAGE on each of four 10% polyacrylamide gels. Following transfer to nitrocellulose membranes and staining with Ponceau-S, nitrocellulose strips containing the fusion proteins were cut, washed with PBS, blocked with 10% dry milk in PBS, washed, and incubated with crude antiserum diluted 1:2 with PBS. Bound antibodies were eluted with 0.1 M glycine, pH 2.5, which was immediately neutralized with 0.1 vols of 1 M Tris, pH 8.0. The buffer was then exchanged to PBS using an Amicon Ultra 15 centrifugal concentrator (Millipore, Bedford, MA), and the purified antibody stored at –80°C.

### Subcellular Fractionation and Western Blots

Neuro2a cells were fractionated essentially by the method of de Duve et al. [21] as previously described [16] to obtain fractions enriched in nuclei (N), mitochondria (M), peroxisomes (L), microsomes (P), and cytosol (S). Proteins were separated by SDS-PAGE and transferred to nitrocellulose membranes. For analysis of subcellular fractions, blots were incubated for 1 hr using blocking buffer (Rockland Immunochemicals, Rockville, MD) and then overnight with primary antibody in a 1:1 solution of blocking buffer and PBS. After washing and incubation with secondary antibodies conjugated to either Alexa fluor 680 or IR dye 800, detection was done using a LiCor Odyssey instrument and software (Lincoln, NE). For detection of ACSF2 overexpressed in COS-1 cells, blots were blocked overnight using 10% dry milk in Tris-buffered saline containing 0.5% Tween. Membranes were incubated with primary and horseradish peroxidase-conjugated secondary antibodies in the same solution, and enhanced chemiluminescence detection was used for detection (SuperSignal West Pico, Pierce).

### Immunohistochemistry and indirect immunofluorescence

Tissues from 3 month old mice were harvested, quickly frozen in liquid nitrogen, and stored at - 80°C. Tissue sections were cut using a cryostat and fixed with 4% paraformaldehyde as described previously [22]. Brain sections were 20 μm thick. Sections of liver, adrenal gland, testis, and ovary were 5-8 μm thick. After fixation, sections were incubated for 30 min with 0.6% H_2_O_2_ in methanol followed by 20 min in 5% normal goat serum. Sections were then incubated sequentially with Avidin D and biotin for 15 min each (Avidin/Biotin blocking kit, Vector Laboratories). Incubation with primary rabbit antibody for ACSF2 was 1 hr at 37°C or overnight at 4°C. Peroxidase based detection was done using a Vectastain ABC kit (Vector laboratories). After the counterstaining with hematoxylin - Harris stain for 30 sec, sections were dehydrated and mounted with DPX - mounting solution (Fluka Biochemika). Indirect immunofluorescence assays of cells plated on glass coverslips were done as described [23].

## RESULTS

### Identification of ACSF2

*ACSF2* was first discovered as part of our effort to identify all candidate ACS genes in the human genome [2] and to characterize the functional roles of their protein products in the brain. Human ACSF2 contains five highly conserved amino acid sequence motifs that are characteristic of all previously reported ACSs [2]. We utilized a conserved 36-37 amino sequence referred to as “Motif II” to segregate ACSs into subfamilies that preferred short-, medium-, long-, very long-chain fatty acid substrates, and the “bubblegum” subfamily. However, four candidate ACSs (including ACSF2) could not be categorized by this criterion, and were designated ACSF (for “ACS family”) members 1-4. Homologs of the ACSF2 gene are present in numerous species of mammal, bird, fish, reptile, insect, and nematode (http://www.ncbi.nlm.nih.gov). Human, mouse, and rat ACSF2 genes encode proteins that contains 615-640 amino acids with a calculated molecular weight of ∼62 kDa. No organellar targeting signals or endoplasmic reticulum protein retention signal were detected (PSORT II; http://psort.ims.u-tokyo.ac.jp/). The protein is predicted to have one transmembrane spanning domain (PredictProtein; http://www.predictprotein.org/).

### Tissue and Cellular Distribution of ACSF2

To determine the expression pattern of ACSF2, Northern blot analysis was done using RNA isolated from different mouse tissues. As shown in Fig. 1A, ACSF2 was expressed in most tissues analyzed except skeletal muscle and perhaps thymus. It is highly expressed in liver, intestine, adrenal gland and ovary, with somewhat weaker expression in brain, heart, kidney, stomach, and testis. To investigate a potential ACSF2 role in brain function, we performed Northern blot analysis on RNA isolated from embryonic and newborn mouse brain. As shown in Fig. 1B, ACSF2 was expressed in brain tissue from embryonic day 12.5 through postnatal day 7; lower expression was found in adult brain.

**Figure 1.**
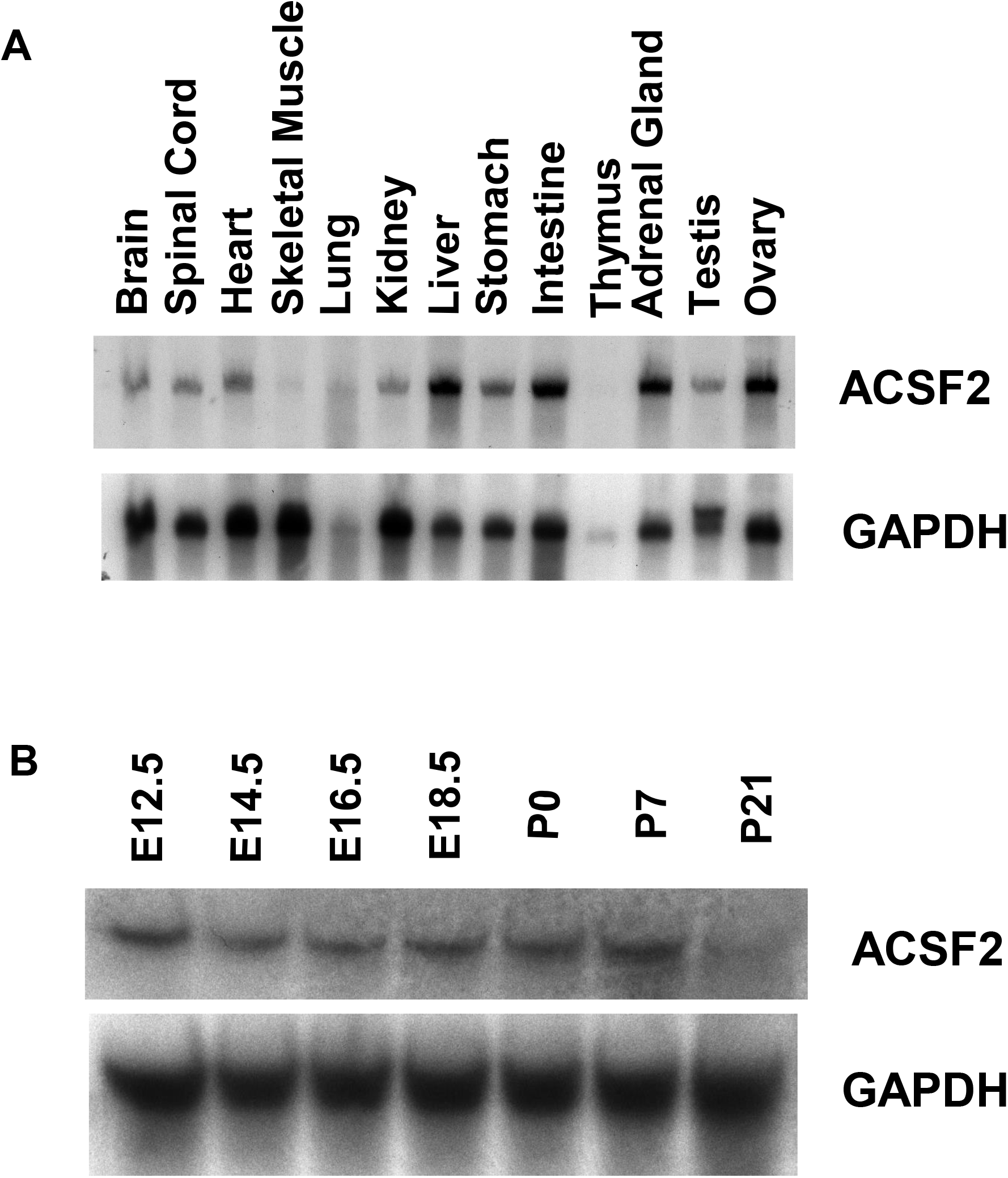
Northern blot analysis. A) Total RNA was isolated from various mouse tissues, separated by agarose gel electrophoresis, transferred to a nitrocellulose membrane, and the membrane incubated with a labeled probe for either ACSF2 (upper panel) or GADPH (lower panel). B) Total RNA was isolated from embryonic (E) and postnatal (P) mouse brain (days E12.5 through P21). Northern blot analysis was performed as in A.

To identify specific cell types that express ACSF2, immunohistochemistry of several different tissues was performed using affinity-purified ACSF2 antibody. In the brain, ACSF2 staining was seen mainly in neurons, including cortical neurons (Fig. 2A), hippocampal neurons (Fig. 2B) and cerebellar Purkinje cells (Fig. 2C). There was no significant staining of glial cells. ACSF2 was ubiquitously expressed in liver, primarily in hepatocytes (Fig. 2D). Within the kidney, strong ACSF2 staining was observed in the cortex (Fig. 2E). There was strong staining of proximal tubular cells but weaker staining of distal tubules; collecting ducts did not express ACSF2. While glomerular cells showed very little ACSF2 expression, intense staining of juxtaglomerular cells and macula densa cells was seen (Fig. 2F). In the adrenal gland, ACSF2 was strongly expressed in all three zones of the cortex (Fig. 2G). Zona glomerulosa cells stained somewhat less intensely than zona fasciculata or zona reticularis cells. ACSF2 was not expressed in the adrenal medulla. In ovary, ACSF2 was ubiquitously expressed (Fig. 2H). Expression was highest in theca cells with weaker staining in the follicle cells. Expression of ACSF2 in testis was found mainly in the interstitial Leydig cells (Fig. 2I). ACSF2 immunostaining was also seen in Sertoli cells, but little staining of spermatocytes was observed.

**Figure 2.**
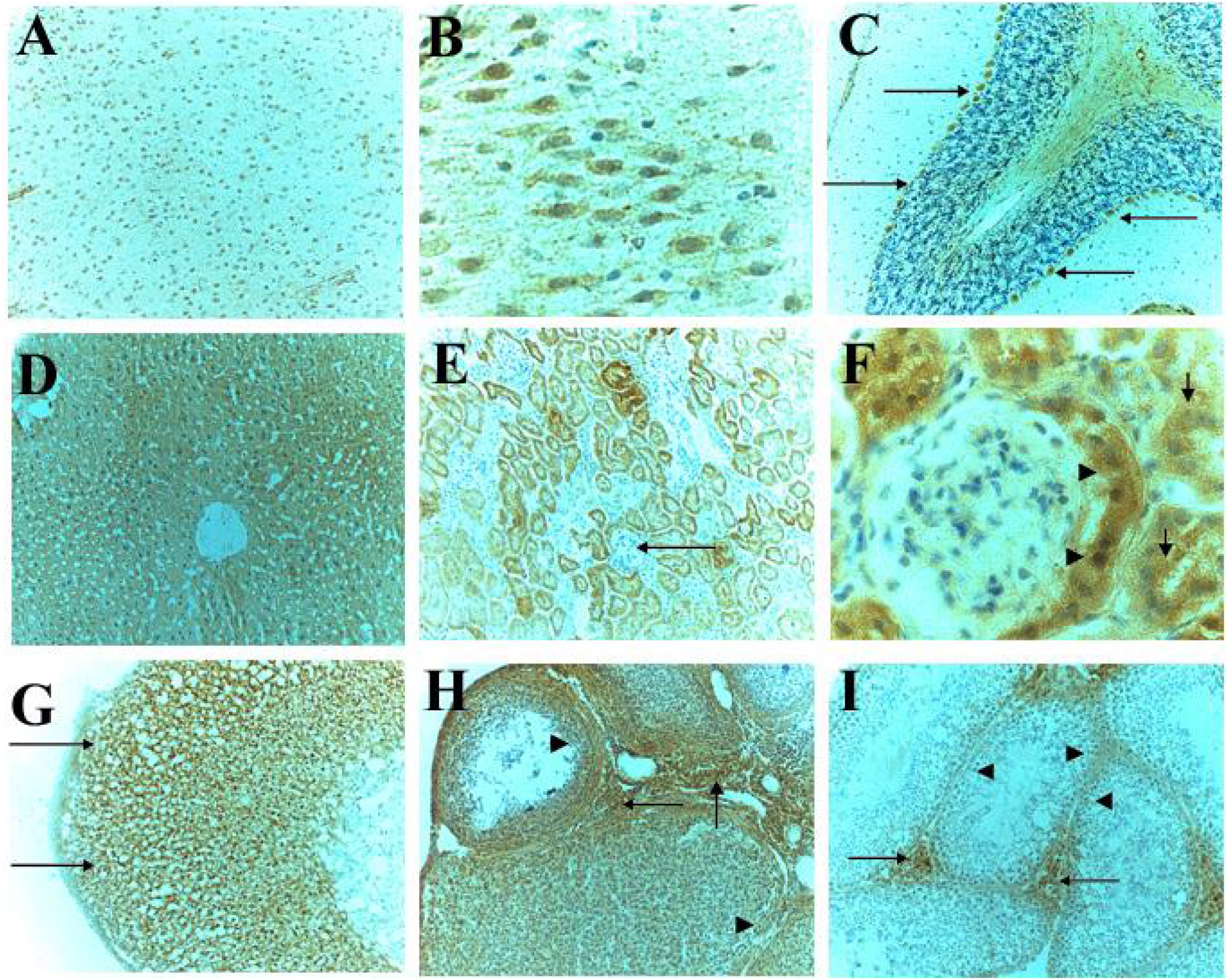
Immunolocalization of ACSF2 in mouse tissues. Tissues were harvested from 3 month old mice and immediately frozen. Preparation of frozen sections and immunohistochemical staining for ACSF2 (brown) was done as described in Experimental Procedures. Sections were counterstained with hematoxylin (blue). A-C, brain. A, temporal cortex; B, hippocampus; C, cerebellum. Staining was mainly observed in neurons, including cerebellar Purkinje cells (arrows). D, liver, showing strong ACSF2 expression in hepatocytes. E and F, kidney. In the renal cortex (E), intense expression was seen in proximal tubule cells, with weaker staining in distal tubules. While glomeruli (E, arrow, and F) did not express ACSF2, juxtaglomerular cells (arrows) and macula densa cells (arrowheads) had a high level of expression. G, adrenal gland, revealing ACSF2 immunostaining in the cortex (arrow) but not the medulla (arrowhead). Expression in the thin, outer layer of adrenal cortex (zona glomerulosa) was somewhat lower than in the broad middle layer (zona fasciculata) and thin inner region (zona reticularis). H, ovary, showing strong ACSF2 expression in theca cells (arrows) and somewhat weaker staining in follicular cells (arrowheads). I, testis, showing robust immunostaining of Leydig cells (arrows) and weaker staining of Sertoli cells (arrowheads).

### Subcellular location of ACSF2

Because immunohistochemical studies showed that ACSF2 is expressed in neurons, we studied mouse Neuro2a neuroblastoma cells to investigate the properties of the endogenous protein. To determine the subcellular location of ACSF2, Neuro2a cells were homogenized and subjected to differential centrifugation. Western blotting was done as described in Experimental Procedures. ACSF2 sedimented in the mitochondria-enriched M-fraction, as shown in Fig. 3 (upper panel). ACSF2 was not found in the N- (nuclear), L- (peroxisome-enriched), P- (microsomal), and S- (cytosolic) fractions. To localize ACSF2 more precisely, Neuro2a cells were analyzed by indirect immunofluorescence using ACSF2 antibody. As demonstrated in Fig. 4A and D, ACSF2 immunostaining was typically found in a single peri-nuclear region in each cell, suggesting a Golgi localization. Double staining with the Golgi-specific markers Golgin-97 (Fig. 4B and C) and GM130 (Fig. 4E and F) demonstrated significant, but incomplete, colocalization with ACSF2.

**Figure 3.**
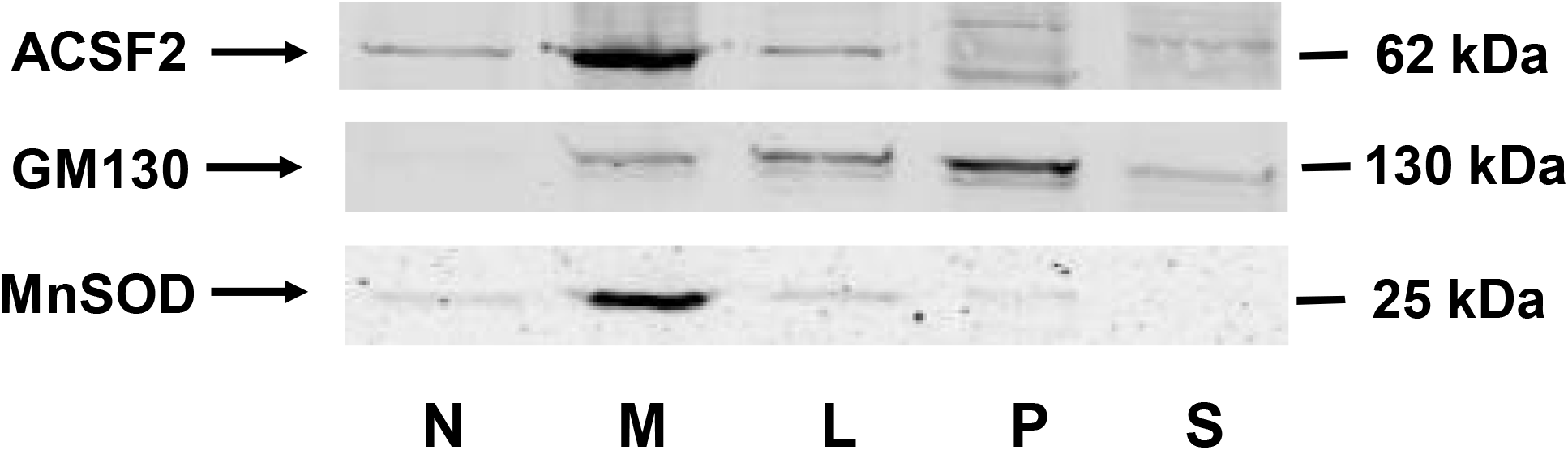
Subcellular localization of ACSF2. Neuro2a cells were homogenized and fractionated by differential centrifugation as described in Experimental Procedures. The resulting subcellular fractions were run on 10% SDS-PAGE and transferred to a nitrocellulose membrane. The membrane was incubated with affinity-purified ACSF2 antibody and detected with using chemiluminescence (upper panel). The membrane was stripped and reprobed using antibody to the Golgi protein GM130 (middle panel). The membrane was stripped again and reprobed using antibody to the mitochondrial protein MnSOD (lower panel).

**Figure 4.**
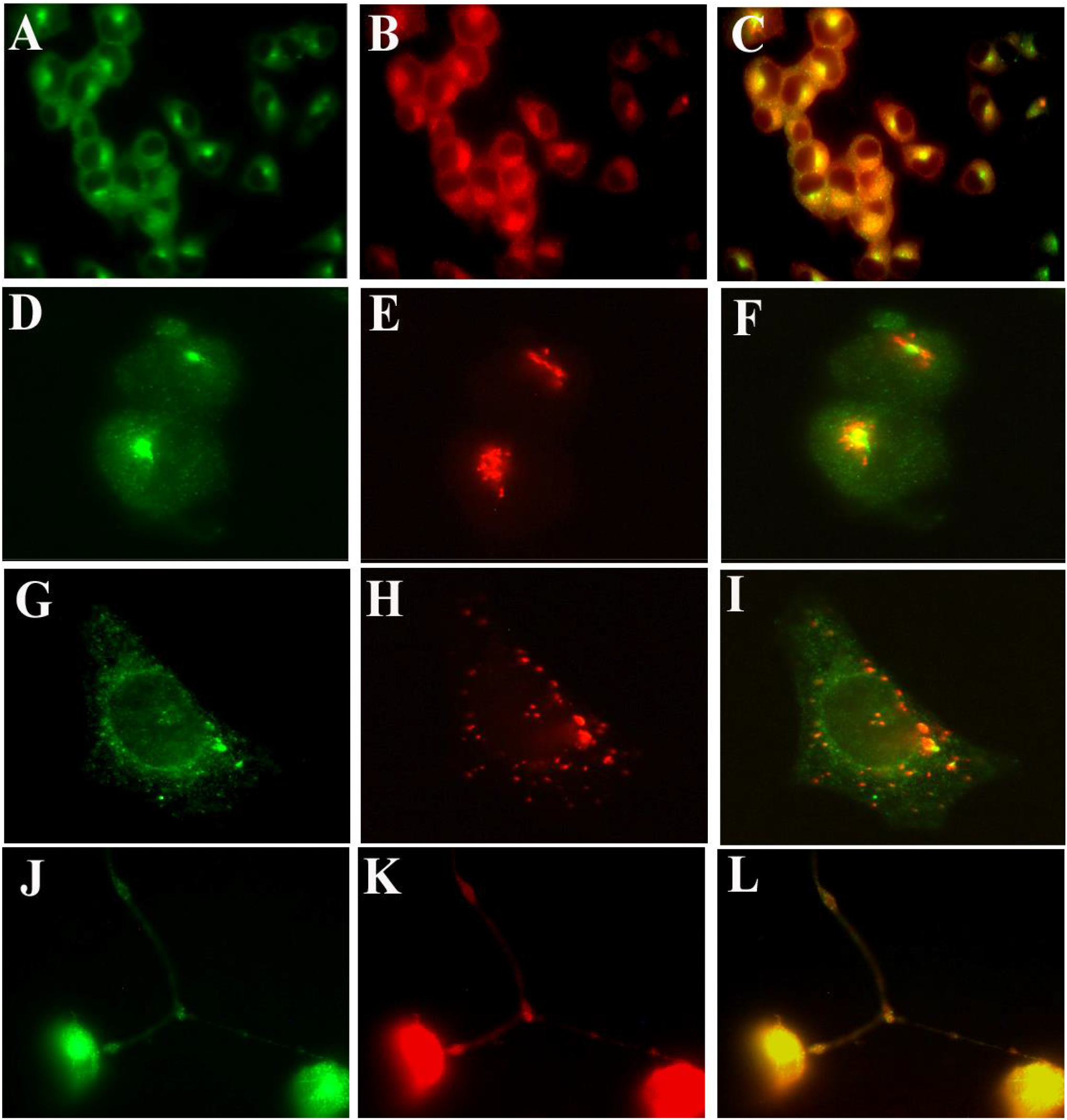
ACSF2 localizes to the Golgi in Neuro2a cells and migrates to synaptic nodes upon differentiation. Neuro2a cells were cultured on glass coverslips, fixed, and permeabilized with Triton X-100. ACSF2 immunostaining (left-hand column of images) was done using affinity-purified rabbit polyclonal antibody and detection was with FITC-conjugated donkey anti-rabbit IgG. Antibodies used for colocalization (middle column of images) were all mouse monoclonals and were detected with rhodamine-conjugated donkey anti-mouse secondary antibodies. Merged images are shown in the right-hand column. A-F. Untreated Neuro2a cells. A. ACSF2; B. Golgin 97; C. merged image; D. ACSF2; E. GM130; F. merged image. G-I. Neuro2a cells treated for 2 hrs with 20 μM nocodazole; G. ACSF2; H, GM130; I. merged image. J-L. Neuro2a cells treated for 2 days with 40 μM retinoic acid; J. ACSF2; K. synaptophysin; L. merged image. A-C, low magnification (200x); D-L, high magnification (1000x).

To confirm this observation, the Western blot shown in Fig. 3 was stripped and reprobed with anti-GM130 (Fig. 3, middle panel). Curiously, this Golgi marker was detected mainly in the P fraction, with smaller amounts of GM130 found in M, L and S fractions. As a control to verify that the M fraction indeed contained mitochondrial proteins, the blot was stripped and probed with antibody for Mn-superoxide dismutase (Fig. 3, lower panel). Like ACSF2, mitochondrial Mn-superoxide dismutase was also found mainly in the M-fraction.

Because of the inconsistency between the Western blot and immunofluorescence results, we explored colocalization of ACSF2 with the Golgi further by treating Neuro2a cells with nocodazole, a microtubule assembly inhibitor which disrupts the Golgi apparatus. Two hours after treatment, the staining patterns of both GM130 and ACSF2 were dramatically different (Fig. 4 G-I). ACSF2 staining was more diffuse throughout the cytoplasm, but a small area of more intense staining remained (Fig. 4G). The GM130 antibody detected numerous punctate vesicles that were also distributed throughout the cytoplasm (Fig. 4H). However, following Golgi disruption, ACSF2 and GM130 no longer colocalized (Fig. 4I). The results indicate that ACSF2 likely resides in the Golgi apparatus in Neuro2a cells, but not primarily in the *cis*-Golgi.

We next used immunofluorescence to determine whether the Golgi localization for ACSF2 was also found in other cell types. Like Neuro2a cells, ACSF2 had a Golgi specific localization in mouse P19 teratocarcinoma cells, as indicated by colocalization with Golgin 97 (Fig. 5A-C); both Neuro2a cells and P19 cells have been studied as models for neurogenesis [24]. In contrast to Neuro2a and P19 cells, however, mouse MA10 (testicular Leydig) cells (Fig. 5D-F), human HepG2 (hepatoma) cells (Fig. 5G-I), and mouse skin fibroblasts (Fig. 5J-L) had an ACSF2 immunostaining pattern more consistent with mitochondria than Golgi, as indicated by colocalization with ATP synthase. A mouse glial cell line, A787, showed minimal ACSF2 immunofluorescent staining (data not shown), in agreement with the results of immunohistochemical staining shown in Fig. 2A-C.

**Figure 5.**
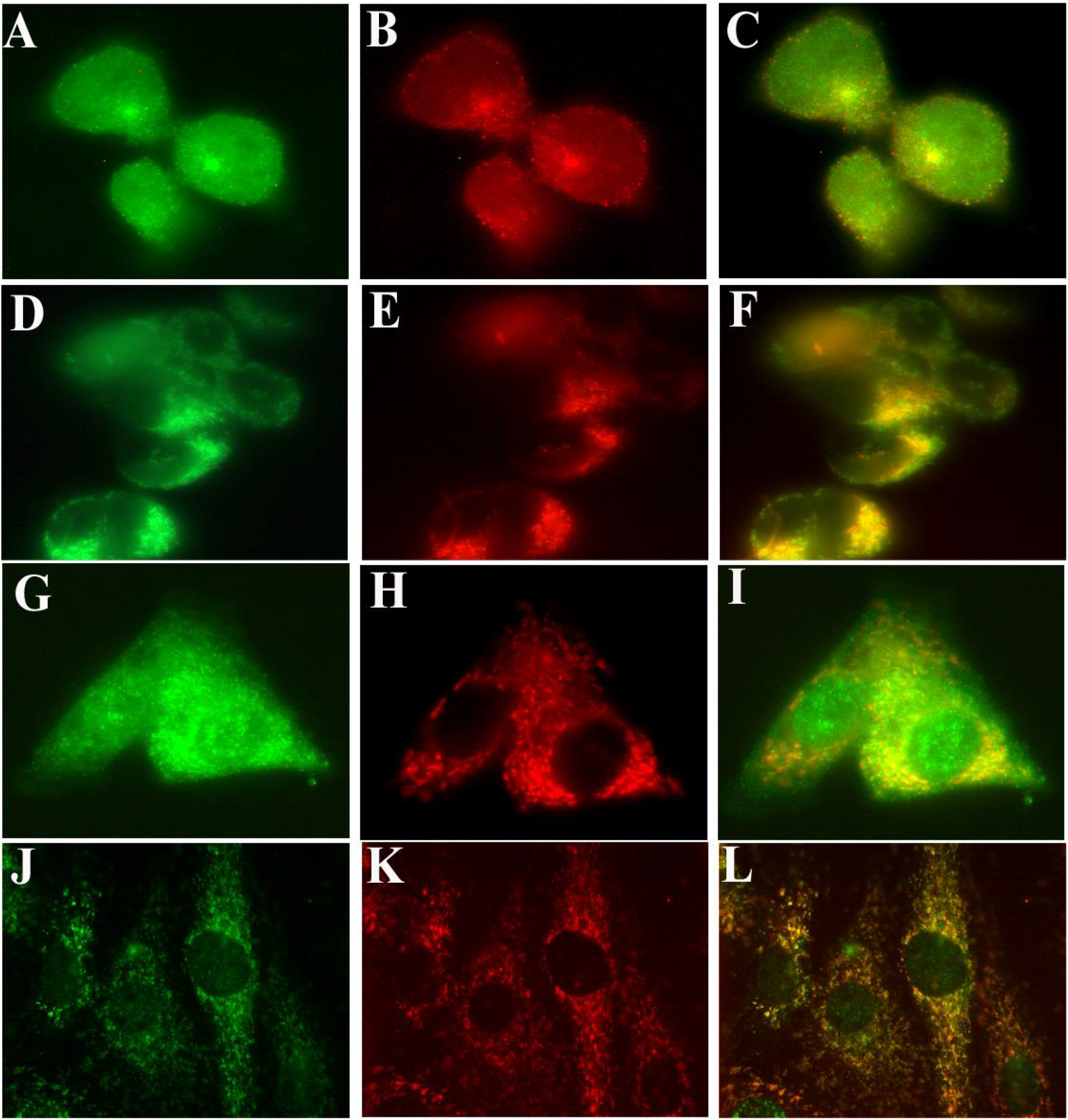
ACSF2 subcellular localization in different cell types. Cells were cultured on glass coverslips, fixed, and permeabilized with Triton X-100. ACSF2 immunostaining (left-hand column of images) was done using affinity-purified rabbit polyclonal antibody and detection was with FITC-conjugated donkey anti-rabbit IgG. Antibodies used for colocalization (middle column of images) were mouse monoclonals and were detected with rhodamine-conjugated donkey anti-mouse secondary antibodies. Merged images are shown in the right-hand column. A-C. P19 cells; A. ACSF2; B. Golgin 97; C. merged image. D-F. MA-10 cells; D. ACSF2; E. ATP synthase; F. merged image. G-I. HepG2 cells; G. ASCF2; H. ATP synthase; I. merged image. J-L. Fibroblasts; J, ACSF2; K. ATP synthase; L. merged image. All panels were high magnification (1000x).

### ACS Activity of mouse brain ACSF2

To examine the enzymatic properties of ACSF2, two approaches were used – overexpression of exogenous protein in COS-1 cells, and siRNA-mediated knockdown of the endogenous protein in Neuro2a cells. We previously categorized ACSF2 as a medium-chain ACS when the enzyme was overexpressed in COS-1 cells, which have low background ACS enzyme activity [2]. Cell extracts were assayed with a representative medium-chain (C8:0), long-chain (C16:0), and very long-chain (C24:0) fatty acid substrate, and only C8:0 activation was significantly increased in cells expressing ACSF2 [2]. To extend this observation, we assayed ACSF2-overexpressing cells with ^14^C-labeled fatty acids of chain lengths ranging from 6-24 carbons. As shown in Fig. 6A, ACSF2 robustly activated medium-chain FA containing 6-10 carbons. The enzyme was weakly active with C12:0, and activity with C14:0 was barely detectable. C16:0, C18:0, C18:1, C18:2, C24:0 fatty acids were also tested as substrates, but no activation was observed (data not shown). Based on these data, ACSF2 can be classified functionally as a medium-chain ACS. Using C10:0 as a substrate, we determined the Km_app_ and Vmax_app_ using the COS-1 cellular extracts overexpressing ACSF2. The Km_app_ was found to be 24.4 μM and the Vmax_app_ was determined to be 385 nmol/20min/mg (Fig. 6B).

**Figure 6.**
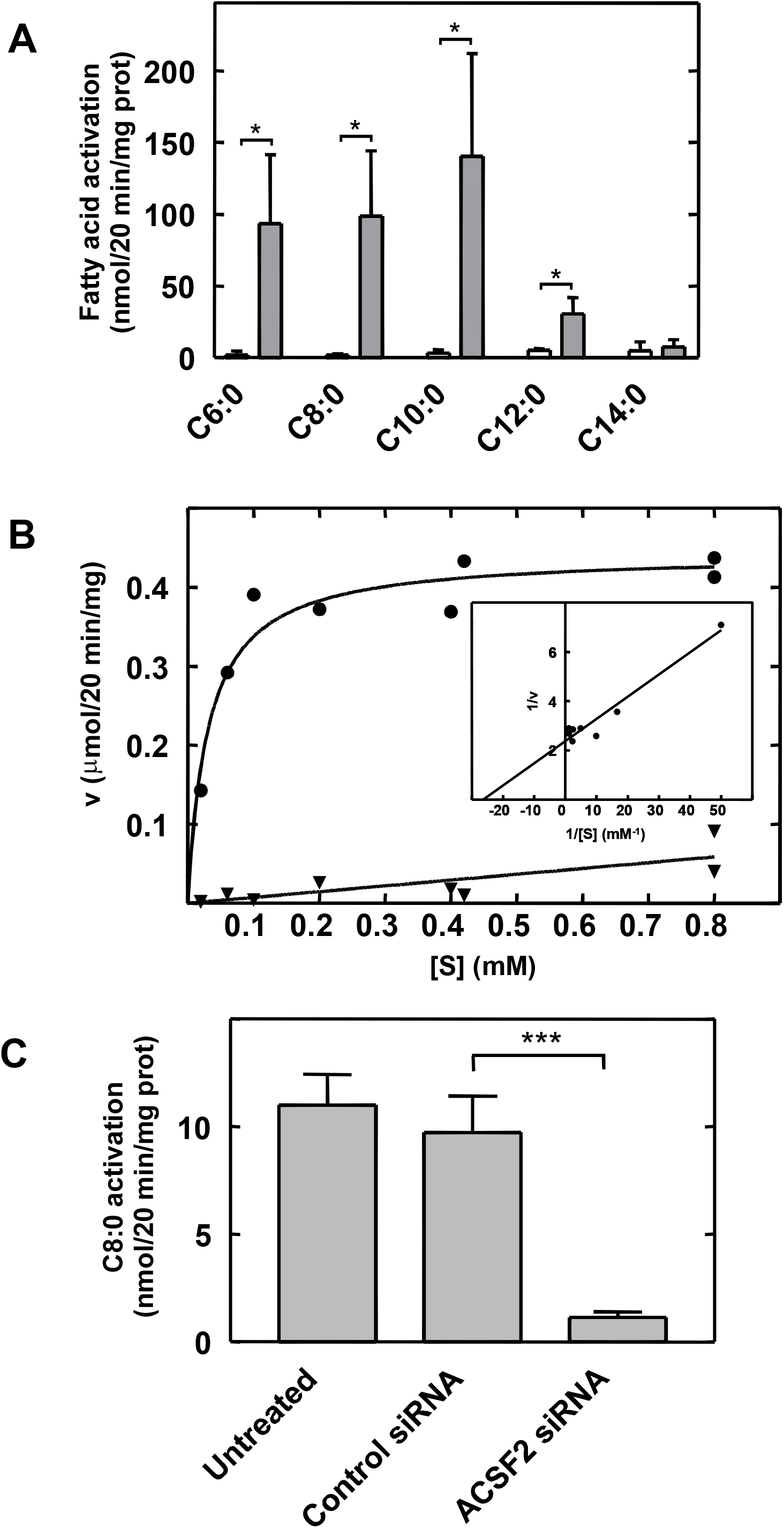
ACS activity of ACSF2. A. COS-I cells were transfected with either full-length ACSF2 in pcDNA3 or empty vector as described in Experimental Procedures. Frozen-thawed cell suspensions (10 μg) were assayed for their ability to activate saturated [1-^14^C]fatty acids (20 μM) containing 6 to 14 carbons. White bars, vector-transfected cells; gray bars, ACSF2-transfected cells. Results shown are mean ± standard deviation of 3 independent transfections. B. Kinetics of ACSF2 activation of C8:0. COS-1 cell suspensions (10 μg) overexpressing ACSF2 were assayed with increasing concentrations of C8:0. After subtraction of the activity measured at each concentration with vector-transfected cells, Michaelis-Menton and Lineweaver-Burk (inset) plots were used to determine Km_app_ and Vmax_app_. C. Endogenous ACSF2 activity of mouse neuroblastoma cells. Neuro2a cells were transfected with either siRNA specific for ACSF2 or control siRNA and harvested three days later. Frozen-thawed cell suspensions (70 μg) were incubated with 400 μM C8:0 as described in Experimental Procedures. Results shown are mean ± standard deviation of 3 independent transfections. Statistical significance was assessed using Student’s t-test (*, p<0.05; ***, p<0.001).

In contrast to the Golgi or mitochondrial subcellular location of endogenous ACSF2 in various cell types (Fig. 5), this protein was found in a diffuse cytoplasmic pattern when overexpressed in COS-1 cells (data not shown). Previous studies with other ACSs revealed that the substrate preference of an endogenous enzyme may differ from that of the overexpressed protein when the latter is mistargeted [16, 25]. To determine activity of endogenous ACSF2, we first measured the capacity of Neuro2a cells to activate [1-^14^C]C8:0 (Fig. 6C). To determine whether ACSF2 plays a significant role in C8:0 metabolism in these cells, we also transiently knocked down ACSF2 protein levels by RNA interference. As shown in Fig. 6C, when Neuro2a cells were treated with ACSF2-specific small interfering double-stranded RNA (siRNA) molecules, C8:0 activation in Neuro2a cells was reduced by at least 90%. Treatment with a non-specific siRNA construct did not significantly decrease the ability of Neuro2a cells to activate C8:0. These results indicate that ACSF2 is responsible for most of the endogenous C8:0 activation in Neuro2a cells.

### Lysine 599 is critical for mouse ACSF2 enzyme activity

All ACS reactions are thought to proceed by a Bi Uni Uni Bi Ping-Pong mechanism [26]. In the ATP-dependent first half-reaction, a fatty acyl substrate is adenylated, with release of inorganic pyrophosphate. Formation of the fatty acid-CoA thioester bond in the second half-reaction releases AMP. It was reported that in a short-chain ACS from *Salmonella enterica*, propionyl-CoA synthetase (PrpE), lysine 592 was required for formation of propionyl-CoA in the first half-reaction [27]. This lysine is found within the conserved 10-amino acid sequence we referred to as Motif V [2]. As illustrated in Fig. 7A, the Motif V amino acid sequences of ACSF2 and PrpE are similar. To test whether ACS activity of mouse ACSF2 is also dependent on this lysine residue, we used site-directed mutagenesis to change this residue to alanine; the construct containing the K599A mutation was cloned into a mammalian expression vector and overexpressed in COS-1 cells and Neuro2a cells. Both wild-type ACSF2 and the K599A mutant were expressed robustly (Fig. 7B). COS-1 cells expressing wild-type ACSF2 exhibited strong activation of C10:0 (Fig. 7C). In contrast, ACS activity of the K599A mutant was not significantly greater that that seen in vector-transfected cells (Fig. 7C). Because ACSF2 was found to be the most abundant medium-chain fatty acid activating enzyme in Neuro2a cells, we also expressed wild-type and K599A mutant ACSF2 cDNAs in this cell type. As in COS-1 cells, native ACSF2 but not the K599A mutant was enzymatically active when overexpressed in Neuro2a cells (Fig. 7D).

**Figure 7.**
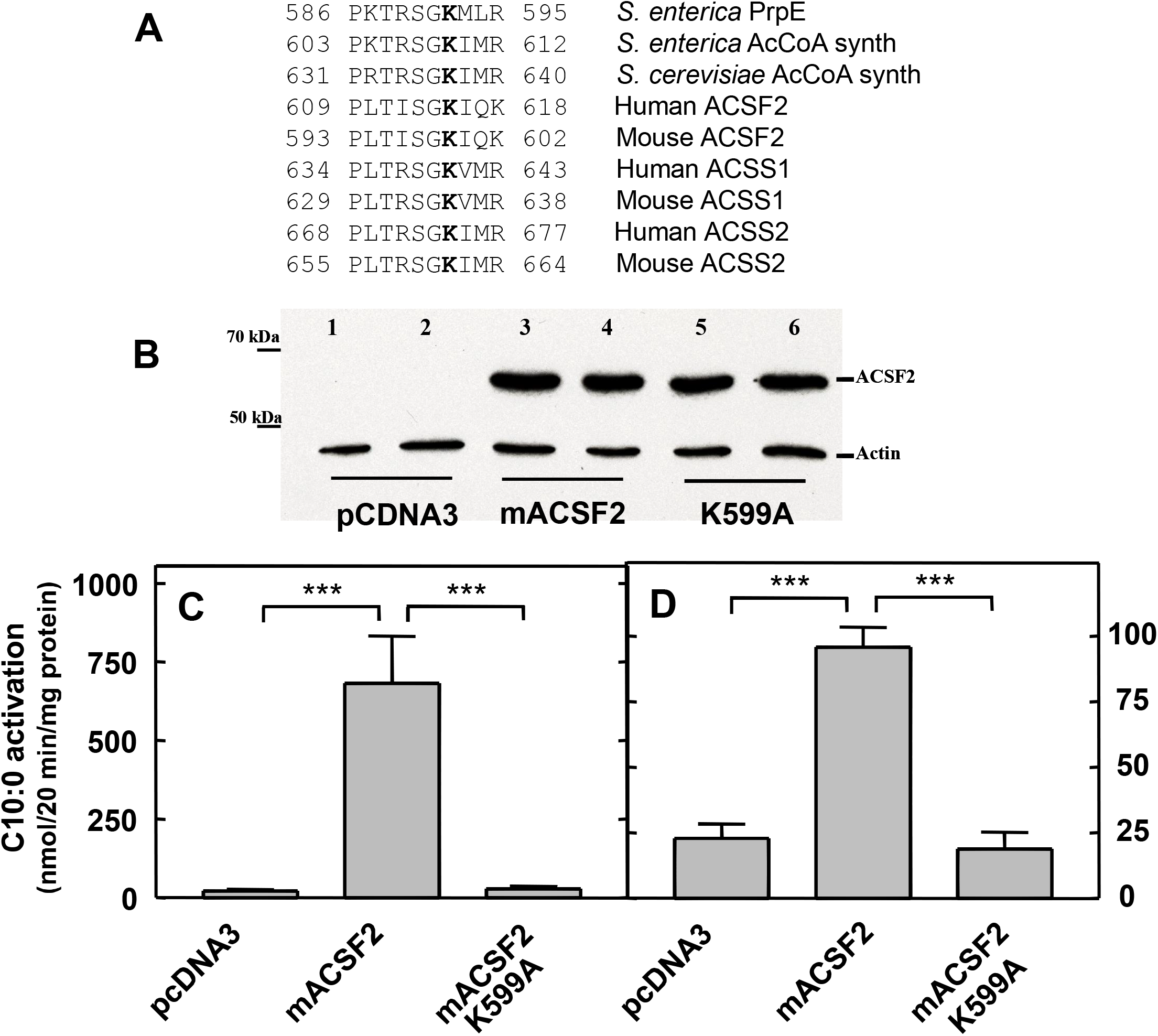
Lys 599 is required for mouse ACSF2 activity. A. Alignment of Motif V amino acid sequences from human and mouse ACSF2 with sequences from *S. enterica* propionyl-CoA synthetase (PrpE), *S. enterica* acetyl-CoA synthetase (Acs), *S. cerevisiae* acetyl-CoA synthetase (Acs2p), human/mouse ACSS1, and human/mouse ACSS2. The conserved lysine, (bold), is residue 599 in mouse ACSF2. B. Wild-type ACSF2 and the K599A mutant are equally expressed in transfected COS-1 cells. Upper panel: Western blot of ACSF2 expression in COS-1 cell suspensions three days post-transfection with pcDNA3 vector (lanes 1 & 2), ACSF2 (lanes 3 & 4), or K599A mutant (lanes 5 & 6). Lower panel: Western blot of the loading control, actin. C & D. ACS activity of wild-type ACSF2 and the K599A mutant in transfected COS-1 cells (C) or transfected Neuro2a cells (D). Cells were harvested and washed, frozen-thawed suspensions (COS-1, 10 μg; Neuro2a, 100 μg) were assayed with 400 μM C10:0. Results shown are mean ± standard deviation of 4 independent transfections. Statistical significance was assessed using Student’s t-test (***, p<0.001)

### Role of ACSF2 in Neuro2A cell lipid metabolism

We next determined whether lack of ACSF2 affected the primary fatty acid utilization pathways in Neuro2a cells. Although the brain does not normally degrade FAs for energy production, it contains an active β-oxidation pathway when assayed with long-chain FA substrates *in vitro*. We assayed β-oxidation of the medium-chain FA C10:0 in the ACSF2 knockdown Neuro2a cell line and control Neuro2a cells expressing normal levels of ACSF2. No differences were observed; control cells degraded the FA at a rate of 0.15±0.03 nmol/hr/mg protein, while knockdown cells did so at a rate of 0.16±0.02 nmol/hr/mg protein. The difference was not statistically significant. It should be noted, however, that β-oxidation activity in these neuronal cells is extremely low. We also incubated control and ACSF2 knockdown Neuro2a cells with [1-^14^C]C10:0 and measured incorporation into neutral lipids and phospholipids after separation by thin-layer chromatography. No specific label incorporation into complex lipids was detectable in either control cells or ACSF2 knockdown cells (data not shown). The results of these studies suggest that medium-chain acyl-CoAs produced by endogenous ACSF2 are not directed towards either degradation or complex lipid synthesis.

### ACSF2 is required for neurite growth in differentiating Neuro2a cells

To determine if ACSF2 plays a role in neuronal differentiation, we treated Neuro2a cells with 40 μM all-*trans*-retinoic acid (RA). After two days exposure to RA, the cells were fixed and stained for ACSF2 and a presynaptic marker, synaptophysin. As shown in Fig. 4J-L, ACSF2 appears to migrate along the developing neurites to synaptic “nodes”.

To define further the role of ACSF2 in developing neurons, we established a Neuro2a cell line expressing a short hairpin RNA (shRNA) construct that stably knocked down the expression of ACSF2. This cell line showed reduced medium chain FA activation (data not shown) similar to that reported in Fig. 6C for Neuro2a cells transiently transfected with siRNA specific for ACSF2. The control cell line was transfected with a plasmid containing a scrambled RNA sequence with no homology to any known gene. After three days of RA treatment, neurite growth was evident in the control cells (Fig. 8A), but appeared to be repressed in the ACSF2-deficient cells (Fig. 8B). To quantitate neurite growth, a pattern of three concentric circles was placed over the cell bodies and the number of concentric circles the longest neurite crossed was counted. Only cells that formed clumps of three cells or less were counted. For ACSF2 knockdown cells, we found that fewer cells developed neurites that crossed one or more concentric circles than for cells transfected with a control shRNA plasmid (Fig. 8C). More than 25% of the control cells developed neurites that crossed one or more circles, as compared to only 3% percent for ACSF2-deficient cells. Reduced neurite length with downexpression of ACSF2 along with constant expression of ACSF2 in developing mouse brain RNA suggests that ACSF2 may play a significant role in brain development.

**Figure 8.**
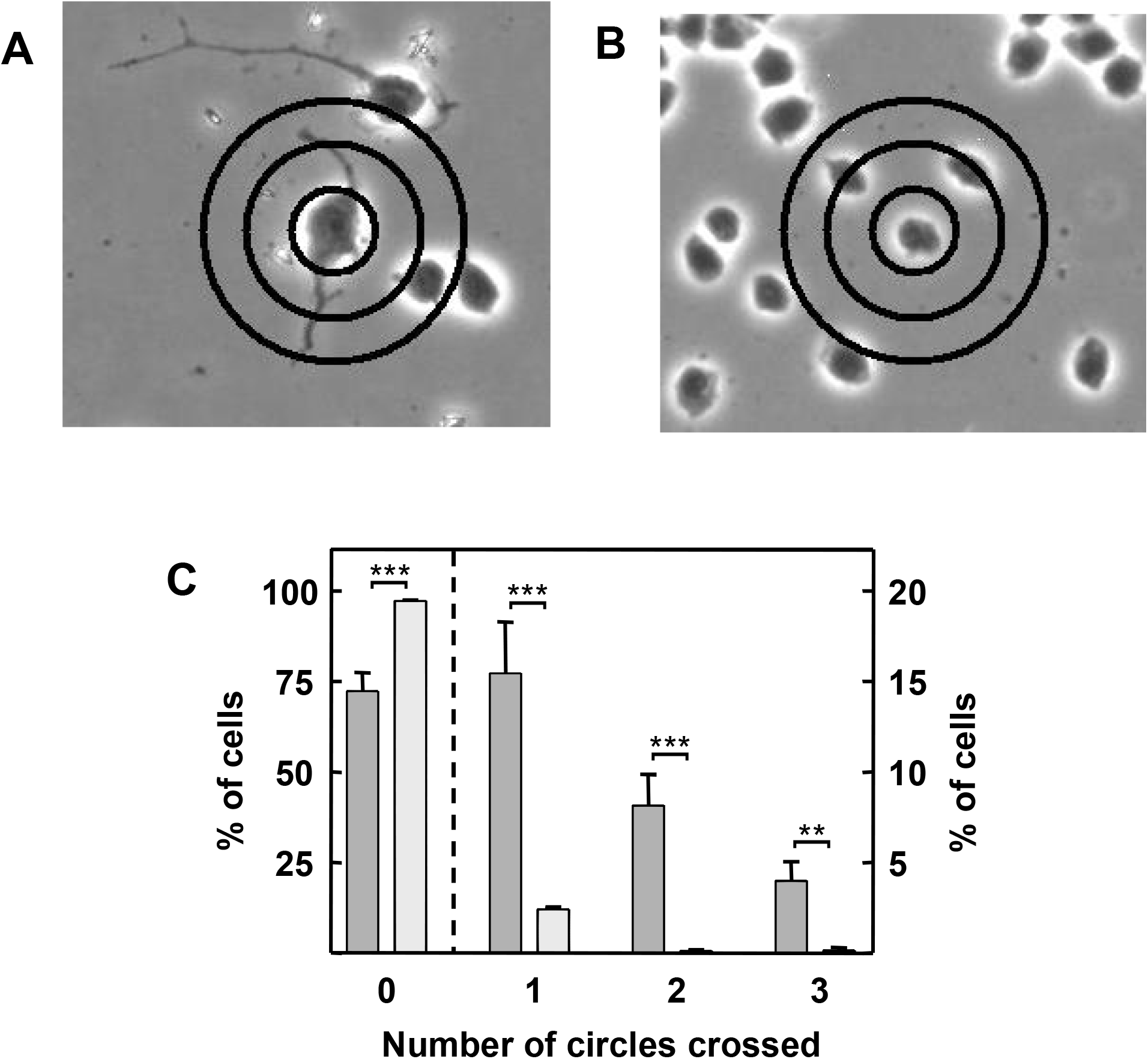
ACSF2 is required for neurite outgrowth in Neuro2a cells differentiated with retinoic acid.. Control and ACSF2-deficient Neuro2a cells as described in the legend to Fig. 6 were treated with 40 μM all-*trans*-retinoic acid. After three days, cells were fixed and imaged at 200x using a phase contrast microscope. A set of three concentric circles were overlaid on the image using Adobe Photoshop and the number of circles the longest axon crossed was tallied as described in Experimental Procedures. A. Control Neuro2a cells. B. ACSF2-deficient Neuro2a cells. C. Percent of cells crossing 0, 1, 2, or 3 circles. Three independent experiments were done. Results are presented as mean ± standard deviation. Approximately 6,000 control and 8,000 ACSF2 knockdown cells were imaged. Statistical significance was assessed using Student’s t-test (**, p<0.01; ***, p≤0.001).

## DISCUSSION

The brain is one of the most lipid-rich organs in the body, and numerous inherited metabolic disorders of the nervous system have at their origin defective enzymes of fatty acid and lipid metabolism. Fatty acids are unable to participate directly in most metabolic pathways unless first activated to their coenzyme A thioesters by enzymes of the ACS family [1,2]. Our interest in neurologic disorders of fatty acid metabolism prompted us to characterize biochemically ACSs of unknown metabolic function, such as SLC27A3 (ACSVL3) [15,28], ACSBG1 [16,25], ACSBG2 [13], and ACSF2 (this work).

Sequence homology of both the entire open reading frame of ACSF2 and conserved motif II suggested that ACSF2 did not readily fit into the short-chain, medium-chain, long-chain, very long-chain, or “bubblegum” subfamilies of ACSs, and phylogenetic analysis indicated that ACSF2 diverged from these subfamilies early in evolution. Thus, we were unable to predict the fatty acyl chain-length preference of ACSF2. Experimentally, ACSF2 appears to be a robust, medium-chain ACS. Its amino acid sequence is slightly more homologous to the six members of the medium-chain (ACSM) subfamily (23-26% amino acid identity) than to all other ACSs (18-22% identity); however, within the ACSM subfamily there is 52-97% identity [2].

Further inspection of the ACSF2 amino acid sequence revealed that it contained a lysine residue (K599 in the mouse protein, and K615 in human ACSF2) in a position previously shown to be critical for the ACS activity of *S. enterica* propionyl-CoA synthetase (PrpE; residue K592) [27]. This lysine residue is conserved in all human ACSs, and is contained within a domain we described as “Motif V” [2]. Mutation of K599 in mACSF2 resulted in nearly complete loss of ACS activity (Fig. 7C, D). The amino acid sequences adjacent to these conserved lysine residues are similar to regulatory domains found in acetyl-CoA synthetases from *S. enterica* and *Saccharomyces cerevisiae* (Fig. 7A). The enzyme activity of the latter proteins are regulated by acetylation/deacetylation of their corresponding lysine residues [29-31]. Additionally, acetylation/deacetylation of homologous lysine residues (Fig. 7A) regulates the enzyme activity of human and mouse acetyl-CoA synthetases ACSS1(formerly AceCS2) and ACSS2 (formerly AceCS1) [32]. The deacetylation of these enzymes is catalyzed by NAD+ dependent enzymes, sirtuins (SIRTs), that affect diverse cellular processes including fatty acid metabolism, apoptosis, and cell cycle [33]. Cytoplasmic SIRT1 regulates cytoplasmic ACSS2, while mitochondrial SIRT3 regulates mitochondrial ACSS1 [32]. Since ACSF2 contains a similar motif, we believe that ACSF2 activity may also be regulated by acetylation/deacetylation of K599. However, since ACSF2 is found in different organelles in different cell types, its regulation may be more complex. The nature of the enzyme or enzymes that regulate ACSF2 activity remains under investigation.

Medium-chain ACSs have been found mainly in mitochondria [34-37]. Although direct experimental evidence is limited, it is generally thought that a primary function of these enzymes is to catalyze the activation of medium-chain fatty acids within the mitochondrial matrix for entry into the β-oxidation pathway. This is a reasonable assumption since medium-chain fatty acids do not require the carnitine shuttle to enter mitochondria. We found that ACSF2 is widely expressed in the liver by immunohistochemistry and is present in the mitochondria of human liver HepG2 cells. Medium-chain fatty acids are abundant in liver from chain shortening of long-chain and very long-chain fatty acids as a result of peroxisomal β-oxidation [38]; these medium-chain fatty acids then migrate to the mitochondria where they are further degraded. However, at least in Neuro2a and teratocarcinoma P19 cells, the subcellular location of ACSF2 was found to be Golgi and not mitochondria, suggesting an alternate metabolic function in these cells.

The high expression of ACSF2 in steroid hormone-producing cells of the adrenal gland, ovary, and testis suggests a potential role in steroidogenesis. In geese, lower ovarian expression of ACSF2 has been associated with increased egg production [39], and ACSF2 polymorphisms were associated with egg laying rate in 3 species of geese [40]. In contrast to geese, egg laying in chickens was positively associated with hepatic ACSF2 expression, and estrogen downregulated ACSF2 [41]. The potential function(s) of ACSF2 in testis and adrenal ACSF2 needs further investigation.

Based on siRNA knockdown of ACSF2 in Neuro2a cells, this enzyme accounts for at least 90% of endogenous medium chain ACS activity (Fig. 6C). Thus, this cell line provides a reasonable model system in which to investigate the metabolic function of ACSF2. As noted above, one potential role for this enzyme is to provide substrate for mitochondrial β-oxidation. We measured the rate of degradation of the medium-chain FA C10:0 by β-oxidation in control and knockdown cells, and found no decrease due to lack of ACSF2. This is consistent with the extramitochondrial subcellular location of ACSF2 in Neuro2a cells. Furthermore, it is unlikely that these cells normally catabolize medium chain FAs since neurons typically use glucose for energy production. Another potential biochemical pathway for utilization of medium-chain fatty acyl-CoA is synthesis of triacylglycerol, as found in lactating mammary gland [42,43]. However, we found no significant incorporation of C10:0 into triacylglycerol, other neutral lipids, or phospholipids in either control or ACSF2 knockdown Neuro2a cells. Thus, the role of ACSF2 in neurons appears to be different than that of a typical medium-chain ACS in liver or mammary gland.

The presence of ACSF2 in Golgi suggested a potential role in protein maturation or protein trafficking. Therefore, we investigated the role of ACSF2 in the differentiation of Neuro2a cells *in vitro* to gain insight into its potential physiologic function. The observation that both ACSF2 and the synaptic vesicle protein, synaptophysin, migrate from the Golgi to regions of potential synapse formation is consistent with a role for ACSF2 in protein or vesicle trafficking. Furthermore, decreased neurite formation in ACSF2-deficient cells suggests that this ACS may play a significant role in synaptogenesis. Synaptophysin is known to migrate from vesicles in the *trans*-Golgi to small synaptic vesicles during the development of neurons [44]. Synaptophysin migration requires an intact microtubule system, the small GTPase, Rab6p [44], and most likely several other vesicle trafficking proteins. Thus, it is possible that lack of ACSF2 interferes with the normal functioning of one of these components. One mechanism by which ACSF2 may contribute to normal vesicle trafficking is by providing activated substrate for protein acylation. Protein acylation with decanoic acid has not been reported, however, and the only protein known to be octanoylated is the orexigenic hormone ghrelin [45]. Ghrelin is produced in stomach and hypothalamus, and its active form is acylated on serine-3 [46]. A potential role for ACSF2 in protein octanoylation or decanoylation is currently under investigation. Lipoic acid is also derived from octanoic acid; when genetically obese (fa/fa) rats were fed lipoic acid, expression of ACSF2 was upregulated, suggesting a potential role for this ACS in lipoic acid metabolism [47].

Several publications have suggested that ACSF2 expression is associated with certain human cancers and ferroptosis. Ferroptosis is a novel oxidative, iron-dependent cell death pathway first described in 2012 [48]. Activation of ferroptosis leads to the non-apoptotic death of cancer cells. The small molecule erastin (eradicator of RAS and ST-expressing cells) is an initiator of ferroptosis, and a screen for erastin resistance genes identified six strong candidates, two of which are lipid metabolism genes ACSF2 and CS (citrate synthase) [48]. Several investigators have used multi-gene screening panels that include ACSF2 (and in some cases, CS), to predict survival in breast cancer [49,50], bladder cancer [51], uveal melanoma [52], and acute myeloid leukemia [53]. Gene fusions are proposed to drive cancer progression, and a MED1-ACSF2 fusion gene has been identified in breast cancer [54]. Further studies are needed to understand the molecular mechanisms underlying ACSF2’s role in cancer.

Finally, more work to understand the physiological function(s) of ACSF2 is needed. In particular, follow-up studies on the disparity between golgi vs. mitochondrial subcellular location are indicated. It is intriguing to speculate that the golgi localization of ACSF2 in cells of neural origin may be important for neurodevelopment. In support of this possibility, Michaelovsky et al. reported a possible association between ACSF2 copy number variation and missense mutations in schizophrenia and neurodevelopment [55].

## ACKNOWLEDGEMENTS

The authors thank Dr. Phyllis Faust (Columbia University) for providing RNA from embryonic and postnatal mouse brain, Dr. Carolyn Machamer (Johns Hopkins University) for expertise and guidance in Golgi localization experiments, and Ms. Meghan L. Maguire for excellent technical assistance. Supported by NIH grants NS37355, HD10981, and HD24061, and the Johns Hopkins University Institutional Research Grant Program.

## ^1^Abbreviations used are

FA: fatty acid
ACS: acyl-CoA synthetase
CoA: coenzyme A
m: mouse
NCBI: National Center for Bioinformatic Information
PBS: phosphate-buffered saline
GAPDH: glyceraldehyde-3-phosphate dehydrogenase

## REFERENCES

1. Watkins PA (1997) Fatty acid activation. Prog Lipid Res. 36:55-83. doi: 10.1016/s0163-7827(97)00004-0.

2. Watkins PA, Maiguel D, Jia Z and Pevsner J (2007) Evidence for 26 distinct acyl-coenzyme A synthetase genes in the human genome. J Lipid Res J Lipid Res. 48:2736–50. doi: 10.1194/jlr.M700378-JLR200.

3. Poulos, A (1995) Very long chain fatty acids in higher animals--a review. Lipids. 30:1–14. doi: 10.1007/BF02537036.

4. Gould SJ, Raymond GV and Valle D (2001) The peroxisome biogenesis disorders. In: The metabolic and molecular bases of inherited disease (Scriver CR, Beaudet AL, Valle D, and Sly WS, eds), 8th Ed., pp. 3181–3217, McGraw Hill, New York.

5. Wanders RJA, Barth PC, and Heymans H A (2001) Single Peroxisomal Enzyme Deficiencies. In: The Metabolic & Molecular Bases of Inherited Disease (Scriver CR, Beaudet AL, Valle D, and Sly WS, eds), Eighth Ed., pp. 3219–3256, McGraw-Hill, New York.

6. Moser HW, Smith KD, Watkins PA, Powers J, and Moser AB (2001) X-linked adrenoleukodystrophy. In: The Metabolic and Molecular Bases of Inherited Disease. (Scriver CR, Beaudet AL, Valle D, and Sly WS, eds), 8th Ed., pp. 3257–3301, McGraw-Hill, New York

7. Mashek DG, Bornfeldt KE, Coleman RA, Berger J, Bernlohr DA, Black P, DiRusso CC, et al. (2004) Revised nomenclature for the mammalian long-chain acyl-CoA synthetase gene family. J Lipid Res. 45:1958–61. doi: 10.1194/jlr.e400002-JLR200.

8. Bowman CE, Rodriguez S, Alpergin ESS, Acoba MG, Zhao L, Hartung T, Claypool SM, Watkins PA, and Wolfgang MJ (2017) The Mammalian Malonyl-CoA Synthetase ACSF3 Is Required for Mitochondrial Protein Malonylation and Metabolic Efficiency. Cell Chem Biol. 24:673–684. doi: 10.1016/j.chembiol.2017.04.009. Epub 2017 May 4.

9. Levtova A, Waters PJ, Buhas D, Lévesque S, Auray-Blais C, Clarke JTR, Laframboise R et al. (2019) Combined malonic and methylmalonic aciduria due to ACSF3 mutations: Benign clinical course in an unselected cohort. J Inherit Metab Dis. 42:107–116. doi: 10.1002/jimd.12032.

10. Lowry OH, Rosebrough NJ, Farr AL, and Randall RJ. (1951) Protein measurement with the Folin phenol reagent. J Biol Chem. 193:265–275.

11. Steinberg SJ, Wang SJ, Kim DG, Mihalik SJ, and Watkins PA (1999) Human very-long-chain acyl-CoA synthetase: cloning, topography, and relevance to branched-chain fatty acid metabolism. Biochem Biophys Res Commun 257:615–621. doi: 10.1006/bbrc.1999.0510.

12. Ho SN, Hunt HD, Horton RM, Pullen JK, and Pease LR (1989) Site-directed mutagenesis by overlap extension using the polymerase chain reaction. Gene 77:51–59. doi: 10.1016/0378-1119(89)90358-2.

13. Pei Z, Jia Z, and Watkins PA (2006) The second member of the human and murine bubblegum family is a testis-and brainstem-specific acyl-CoA synthetase. J Biol Chem. 281(10):6632–41. doi: 10.1074/jbc.M511558200.

14. Ascoli M (1981) Characterization of several clonal lines of cultured Leydig tumor cells: gonadotropin receptors and steroidogenic responses. Endocrinology 108:88–95. doi: 10.1210/endo-108-1-88.

15. Pei Z, Fraisl P, Berger J, Jia Z, Forss-Petter S, and Watkins PA (2004) Mouse very long-chain Acyl-CoA synthetase 3/fatty acid transport protein 3 catalyzes fatty acid activation but not fatty acid transport in MA-10 cells. J Biol Chem. 279:54454–54462. doi: 10.1074/jbc.M410091200.

16. Pei Z, Oey NA, Zuidervaart MM, Jia Z, Li Y, Steinberg SJ, Smith KD, and Watkins PA (2003) The acyl-CoA synthetase “bubblegum” (lipidosin): further characterization and role in neuronal fatty acid beta-oxidation. J Biol Chem 278:47070–47078. doi: 10.1074/jbc.M310075200.

17. Dole VP (1956) A relation between non-esterified fatty acids in plasma and the metabolism of glucose. J Clin Invest. 35:150–154. doi: 10.1172/JCI103259.

18. Fujino T, Kondo J, Ishikawa M, Morikawa K, and Yamamoto TT (2001) Acetyl-CoA Synthetase 2, a Mitochondrial Matrix Enzyme Involved in the Oxidation of Acetate. J Biol Chem 276:11420–11426. doi.org/10.1074/jbc.M008782200.

19. Watkins PA, Ferrell EVJr, Pedersen JI, and Hoefler G (1991) Peroxisomal fatty acid beta-oxidation in HepG2 cells. Arch Biochem Biophys. 289:329–336. doi: 10.1016/0003-9861(91)90419-j.

20. Folch J, Lees M, and Sloane-Stanley GH (1957) A simple method for the isolation and purification of total lipides from animal tissues. J Biol Chem. 226:457–509.

21. de Duve C, Pressman BC, Gianetto R, Wattiaux R, and Appelmans F (1955) Tissue fractionation studies. 6. Intracellular distribution patterns of enzymes in rat-liver tissue. Biochem J. 60:604–617. doi: 10.1042/bj0600604.

22. Mihalik SJ, Steinberg SJ, Pei Z, Park J, Kim DG, Heinzer AK, Dacremont G, Wanders RJ, Cuebas DA, Smith KD, and Watkins PA. (2002) Participation of two members of the very long-chain acyl-CoA synthetase family in bile acid synthesis and recycling. J Biol Chem 277:24771–24779. doi: 10.1074/jbc.M203295200.

23. Watkins PA, Gould SJ, Smith MA, Braiterman LT, Wei H-M, Kok F, Moser AB, Moser HW, and Smith KD (1995) Altered expression of ALDP in X-linked adrenoleukodystrophy. Am J Hum Genet. 57:292–301.

24. Leszczyński P, śmiech M, Salam Teeli A, Haque E, Viger R, Ogawa H, Pierzchała M, and Taniguchi H. (2020) Deletion of the Prdm3 Gene Causes a Neuronal Differentiation Deficiency in P19 Cells. Int J Mol Sci. 21:7192. doi: 10.3390/ijms21197192.

25. Steinberg SJ, Morgenthaler J, Heinzer AK, Smith KD, and Watkins PA. (2000) Very long-chain acyl-CoA synthetases. Human “bubblegum” represents a new family of proteins capable of activating very long-chain fatty acids. J Biol Chem 275:35162–35169. doi: 10.1074/jbc.M006403200.

26. Hisanaga Y, Ago H, Nakagawa N, Hamada K, Ida K, Yamamoto M, Hori T, et al. (2004) Structural basis of the substrate-specific two-step catalysis of long chain fatty acyl-CoA synthetase dimer. J Biol Chem 279:31717–31726. doi: 10.1074/jbc.M400100200.

27. Horswill AR and Escalante-Semerena JC (2002) Characterization of the propionyl-CoA synthetase (PrpE) enzyme of Salmonella enterica: residue Lys592 is required for propionyl-AMP synthesis. Biochemistry 41, 2379–2387. doi: 10.1021/bi015647q.

28. Pei Z, Sun P, Huang P, Lal B, Laterra J and Watkins PA (2009) Acyl-CoA synthetase VL3 knockdown inhibits human glioma cell proliferation and tumorigenicity. Cancer Res 69:9175–9182. doi: 10.1158/0008-5472.CAN-08-4689.

29. Starai VJ, Takahashi H, Boeke JD, and Escalante-Semerena JC (2003) Short-chain fatty acid activation by acyl-coenzyme A synthetases requires SIR2 protein function in Salmonella enterica and Saccharomyces cerevisiae. Genetics 163:545–555. doi: 10.1093/genetics/163.2.545.

30. Starai VJ, Celic I, Cole RN, Boeke JD, and Escalante-Semerena JC (2002) Sir2-dependent activation of acetyl-CoA synthetase by deacetylation of active lysine. Science 298, 2390–2392. doi: 10.1126/science.1077650.

31. Starai VJ and Escalante-Semerena JC (2004) Identification of the protein acetyltransferase (Pat) enzyme that acetylates acetyl-CoA synthetase in Salmonella enterica. J Mol Biol 340:1005–1012. doi: 10.1016/j.jmb.2004.05.010.

32. Hallows WC, Lee S and Denu JM (2006) Sirtuins deacetylate and activate mammalian acetyl-CoA synthetases. Proc Natl Acad Sci U S A. 103:10230–10235. doi: 10.1073/pnas.0604392103.

33. Denu JM (2005) The Sir 2 family of protein deacetylases. Curr Opin Chem Biol 9:431–440. doi: 10.1016/j.cbpa.2005.08.010.

34. Vessey DA, Kelley M and Warren RS (1999) Characterization of the CoA ligases of human liver mitochondria catalyzing the activation of short-and medium-chain fatty acids and xenobiotic carboxylic acids. Biochim Biophys Acta 1428:455–462. doi: 10.1016/s0304-4165(99)00088-4.

35. Kasuya F, Igarashi K and Fukui M (1999) Characterization of a renal medium chain acyl-CoA synthetase responsible for glycine conjugation in mouse kidney mitochondria. Chem Biol Interact 118:233–246. doi: 10.1016/s0009-2797(99)00084-8.

36. Kasuya F, Tatsuki T, Ohta M, Kawai Y and Igarashi K (2005) Purification, characterization, and mass spectrometric sequencing of a medium chain acyl-CoA synthetase from mouse liver mitochondria and comparisons with the homologues of rat and bovine. Protein Expr Purif. 47:405–414. doi: 10.1016/j.pep.2005.11.006.

37. Fujino T, Takei YA, Sone H, Ioka RX, Kamataki A, Magoori K, Takahashi S, Sakai J and Yamamoto TT (2001) Molecular identification and characterization of two medium-chain acyl-CoA synthetases, MACS1 and the Sa gene product. J Biol Chem 276:35961–35966. doi: 10.1074/jbc.M106651200.

38. Hashimoto T (1999) Peroxisomal beta-oxidation enzymes. Neurochem Res 24:551–563. doi: 10.1023/a:1022540030918.

39. Yu S, Wei W, Xia M, Jiang Z, He D, Li Z, Han H et al. (2016) Molecular characterization, alternative splicing and expression analysis of ACSF2 and its correlation with egg-laying performance in geese. Anim Genet. 47:451–462. doi: 10.1111/age.12435.

40. Yang Y, An C, Yao Y, Cao Z, Gu T, Xu Q and Chen G (2019) Intron polymorphisms of MAGI-1 and ACSF2 and effects on their expression in different goose breeds. Gene. 701:82–88. doi: 10.1016/j.gene.2019.02.102.

41. Tian W, Zheng H, Yang L, Li H, Tian Y, Wang Y and Lyu S (2018) Dynamic Expression Profile, Regulatory Mechanism and Correlation with Egg-laying Performance of ACSF Gene Family in Chicken (Gallus gallus). Sci Rep. 8:8457. doi: 10.1038/s41598-018-26903-6.

42. Hachey DL, Silber GH, Wong WW and Garza C (1989) Human lactation. II: Endogenous fatty acid synthesis by the mammary gland. Pediatr Res. 25:63–68. doi: 10.1203/00006450-198901000-00015.

43. Hansen HO, Grunnet I and Knudsen J (1984) Triacylglycerol synthesis in goat mammary gland. The effect of ATP, Mg2+ and glycerol 3-phosphate on the esterification of fatty acids synthesized de novo. Biochem J. 220:513–519. doi: 10.1042/bj2200513.

44. Tixier-Vidal A, Barret A, Picart R, Mayau V, Vogt D, Wiedenmann B and Goud B (1993) The small GTP-binding protein, Rab6p, is associated with both Golgi and post-Golgi synaptophysin-containing membranes during synaptogenesis of hypothalamic neurons in culture. J Cell Sci. 105: 935–947.

45. Davis TR, Pierce MR, Novak SX, Hougland JL (2021) Ghrelin octanoylation by ghrelin O-acyltransferase: protein acylation impacting metabolic and neuroendocrine signaling. Open Biol. 11:210080. doi: 10.1098/rsob.210080.

46. Marzullo P, Verti B, Savia G, Walker GE, Guzzaloni G, Tagliaferri M, Di Blasio A and Liuzzi A. (2004) The relationship between active ghrelin levels and human obesity involves alterations in resting energy expenditure. J Clin Endocrinol Metab. 89:936–939. doi: 10.1210/jc.2003-031328.

47. Pashaj A, Yi X, Xia M, Canny S, Riethoven J-JM, Moreau R (2013) Characterization of genome-wide transcriptional changes in liver and adipose tissues of ZDF (fa/fa) rats fed R-α-lipoic acid by next-generation sequencing. Physiol Genomics. 45:1136–1143. doi: 10.1152/physiolgenomics.00138.2013.

48. Dixon SJ, Lemberg KM, Lamprecht MR, Skouta R, Zaitsev EM, Gleason CE, Patel DN, et al. (2012) Ferroptosis: an iron-dependent form of nonapoptotic cell death. Cell. 149:1060–1072. doi: 10.1016/j.cell.2012.03.042.

49. Wang H, Liu C, Zhao Y and Gao G. (2020) Mitochondria regulation in ferroptosis. Eur J Cell Biol. 99:151058. doi: 10.1016/j.ejcb.2019.151058.

50. Li H, Li L, Xue C, Huang R, Hu A, An X and Shi Y (2021) A Novel Ferroptosis-RelatedGene Signature Predicts Overall Survival of Breast Cancer Patients. Biology. 10, 151. doi.org/10.3390/biology10020151.

51. Zhang S, Wang C, Xia W, Duan H, Qian S, Shen H (2021) Novel Ferroptosis-Related Multigene Prognostic Models for Patients with Bladder Cancer. Int J Gen Med. 14:8651–8666. doi: 10.2147/IJGM.S339996.

52. Jin Y, Wang Z, He D, Zhu Y, Gong L, Xiao M, Chen X and Cao K (2021) Analysis of Ferroptosis-Mediated Modification Patterns and Tumor Immune Microenvironment Characterization in Uveal Melanoma. Front Cell Dev Biol. 9:685120. doi: 10.3389/fcell.2021.685120.

53. Huang R, Liao X and Li Q (2017) Identification and validation of potential prognostic gene biomarkers for predicting survival in patients with acute myeloid leukemia. Onco Targets Ther. 10:5243–5254. doi: 10.2147/OTT.S147717.

54. Zhou L, Song Z, Hu J, Liu L, Hou Y, Zhang X, Yang X and Chen K (2021) ACSS3 represses prostate cancer progression through downregulating lipid droplet-associated protein PLIN3. Theranostics. 11:841–860. doi: 10.7150/thno.49384.

55. Michaelovsky E, Carmel M, Frisch A, Salmon-Divon M, Pasmanik-Chor M, Weizman A and Gothelf D (2019) Risk gene-set and pathways in 22q11.2 deletion-related schizophrenia: a genealogical molecular approach. Transl Psychiatry. 9:15. doi: 10.1038/s41398-018-0354-9.

